# Multi-modal comparison of primary and stem cell-derived β-cells nominates targets for maturation

**DOI:** 10.64898/2026.06.04.730032

**Authors:** Jasmine Maghera, Cara E. Ellis, Aliya F. Spigelman, Nancy Smith, Shugo Sasaki, Francis C. Lynn, Patrick E. MacDonald

## Abstract

Stem-cell-derived β-like-cells (SCβ-cells) provide a promising platform for diabetes modelling and cell replacement therapy, but their incomplete functional maturation remains a challenge. Here, we compared immature SCβ-cells to human primary β-cells utilizing a multi-omic, single-cell framework integrating patch-clamp electrophysiology with scRNA sequencing (patch-seq), regulatory network inference, and functional phenotyping. Despite low insulin secretion and reduced insulin content, SCβ-cells displayed larger Na^+^ and Ca^2+^ currents and depolarization-induced exocytosis. Ultrastructural and metabolic profiling revealed immature insulin granules, altered mitochondrial morphology, elevated basal respiration and proton leak, and diminished spare respiratory capacity and glucose-responsive metabolism. Patch-seq linked exocytotic activity in SCβ-cells to oxidative phosphorylation and MYC target programs, consistent with incomplete terminal differentiation, whereas SCβ-cells expressing higher levels of mature identity markers showed reduced ion channel hyperactivity. Multi-omics profiling showed that electrophysiological features in SCβ-cells were embedded in transcriptional programs distinct from those of primary β-cells and other endocrine cells. Network control theory nominated SREBP1, an endoplasmic reticulum tethered transcription factor regulating cholesterol and lipid homeostasis, as a promising candidate involved in this immature state. Inhibition of cholesterol trafficking increased SREBF1 expression and shifted metabolic and transcriptional features towards a more mature β-like state. These data identify potential targets and pathways that can be leveraged to improve SCβ-cell maturation and validate cholesterol and lipid homeostasis through the SREBP1 axis as one such candidate.

## INTRODUCTION

Type 1 diabetes (T1D) is defined by the destruction of insulin-producing β-cells in the pancreatic islets of Langerhans by the body’s immune system. Transplantation of whole-pancreas or isolated islets has demonstrated long-term benefits.^1^ However, these approaches are limited by strict eligibility criteria, lifelong immunosuppressive therapy, and scarcity of donor organs. Substantial β-cell death immediately post-transplant drives graft attrition, often necessitating transplantation of large numbers of islets from multiple donors and repeated infusions.^1–3^ Stem-cell-derived β-cells (SCβ-cells) offer the potential for an alternative, unlimited supply of implantable material, reduced immunogenicity through genetic modification, and an avenue to model human islet development and disease. Differentiation from human embryonic (hESCs) or induced pluripotent stem cells (hiPSCs) toward a β-cell fate yields cells that are capable of insulin secretion,^4^ express similar canonical gene and protein markers, and can ameliorate diabetes in relevant mouse models^5,6^ and humans.^7^

However, generating SCβ-cells *in vitro* that fully mimic primary β-cells remains an unmet goal. SCβ-cells fail to regulate insulin exocytosis^8,9^ and show a higher reliance on glycolytic rather than oxidative metabolism, with reduced anaplerotic cycling, contributing to impaired glucose sensing.^9–11^ While gene expression profiles of SCβ-cells approach those of primary β-cells during *in vivo* maturation, they still express lower levels of maturation-linked genes such as *MAFA*, *UCN3*, *IAPP,* and *INS* compared to primary β-cells, suggesting the need for further optimization and a more nuanced understanding of transcriptional networks.^5,6,9,12–14^

Single-cell RNA sequencing (scRNA-seq) has provided critical insights into the transcriptional differences between SCβ-cells and primary β-cells, helping to refine differentiation protocols.^5,6,13^ In parallel, electrophysiological techniques such as whole-cell patch-clamp have enabled analysis of functional properties like exocytosis and ion channel dynamics, key determinants of insulin secretion.^6,15^ Integration of scRNA-seq and electrophysiology (patch-seq) and bioinformatic approaches offers a more complete picture of cell physiology by revealing how global transcriptional states drive cellular behavior and function. In this study, we apply a multimodal patch-seq approach^16–18^ and contextualize our findings with functional and structural assays. To characterize transcriptional regulation relative to primary β-cells, we infer gene regulatory networks (GRNs) utilizing single-cell regulatory network inference (SCENIC),^19,20^ and integrate these with patch-seq using supervised pseudobulk and unsupervised single-cell approaches.^21–23^ We find that though SCβ-cells activate some key β-cell programs, these programs exist in a dysregulated context, lacking the modular precision and coordinated electrophysiological function observed in human primary β-cells. Furthermore, some features exhibited by SCβ-cells separate them from primary islet endocrine fates.

Finally, using network control theory, we nominate transcription factors whose network influence and steerability make them compelling candidates for SCβ-cell maturation, and validate a candidate transcription factor, *SREBF1*/SREBP1. Perturbing lipid and cholesterol stores in SCβ-cells increased *SREBF1* expression and that of β-cell identity markers, while decreasing immaturity transcripts and improving oxygen consumption profiles. These data support the idea that achieving mature β-cell function requires a broad rewiring of SCβ-cell regulatory and functional modalities and identify cholesterol homeostasis through the SREBP1 axis as one tractable entry point.

## RESULTS

### Immature SCβ-cell function despite robust ion channel activity and exocytosis

We studied SCβ-cells at two different stages: nascent SCβ-cells at day 30 of culture (SCβ D30) and cells further cultured to approximately day 50 (SCβ D50), compared to primary islets and β-cells from non-diabetic donors (**Fig. 1A**). Like primary islets, SCβ-cell clusters with INS-driven eGFP expression promoter^24,25^ stained positive with dithizone, indicating the likely presence of Zn^2+^-containing insulin granules (**Fig. 1B**). Cell clusters were assessed for static and dynamic glucose-stimulated insulin secretion (GSIS), metabolic analysis, or dissociated into single cells to assess depolarization-induced exocytosis,Na^+^ and Ca^2+^ currents, and other electrophysiological parameters by whole cell patch-clamp. Static glucose-stimulated insulin secretion (GSIS) was much lower in SCβ D30 cells and SCβ D50 cells compared to primary islets, with median total insulin contents being 2.31 and 3.80-fold lower, respectively (**Fig.1C**). Consistent with the static GSIS results, a dynamic perifusion assay (**Fig. 1D**) similarly showed little response of the SCβ-cells to increased (16.7 mM) glucose, however, some KCl-induced insulin secretion was observed. Although others have reported glucose-dependent insulin secretion from SCβ-cells,^6,9^ these findings are consistent with an immature state of the SCβ-cells studied here and by others^13,26^. As such, further analysis focused on understanding differences between immature SCβ-cells and primary β-cells.

**Figure 1:**
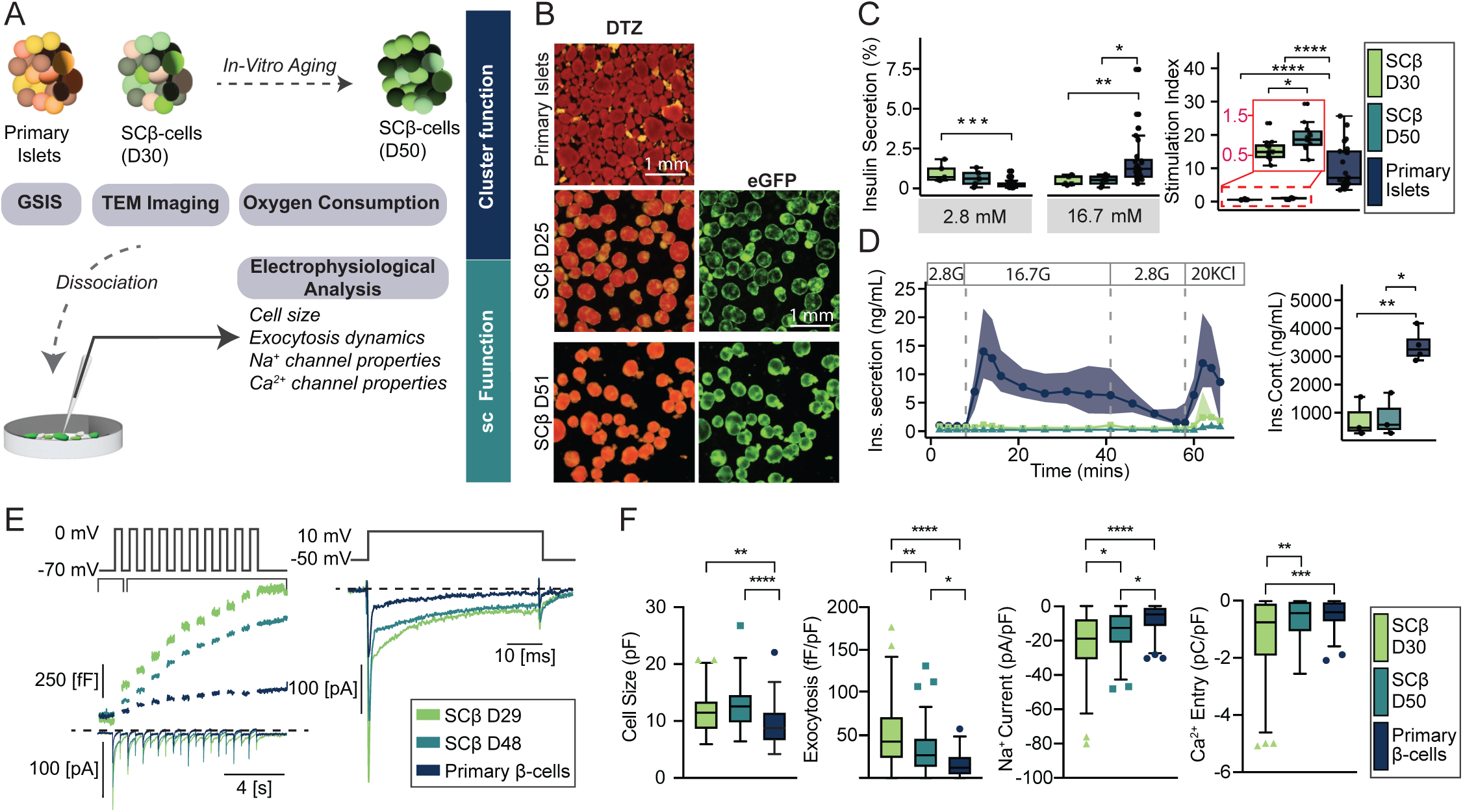
Immature SCβ-cell function is characterized by aberrant electrophysiological profiles and dysregulated Insulin Secretion. (A) Schematic workflow illustrating the experimental design, including *in vitro* SCβ-cell differentiation, aging to approximate day 30 (D30) and day 50 (D50), and downstream functional assessments. (B) Representative images of dithizone (DTZ) staining and eGFP fluorescence in primary human islets and SC&-cell clusters at D25 and D51. Scale bars, 1 mm. (C) Static insulin secretion responses in D30 and D50 SCβ-cells measured at 2.8 mM glucose and 16.7 mM glucose. Responses were normalized to total insulin content. N=5 for D30, N=5 for D50 SCβ-cells and N=24 for primary human islets. (D) Dynamic insulin secretion responses in SCβ-cells at D30 and D50 measured during sequential stimulation with 2.8 mM glucose, 16.7 mM glucose, and 2.8 mM glucose + 20 mM KCl. N=3 for D30, N=3 for D50 SCβ-cells and N=5 for primary human islets. (E) Representative electrophysiology traces showing changes in membrane capacitance (fF) and ionic current (pA) in SCβ-cells and primary β-cells during a train of 10 membrane depolarizations (*left*) or a single 500 ms depolarization (*right*). (F) Quantification of electrophysiological parameters across cell types, including cell size (as initial membrane capacitance; pF), and total exocytosis responses (fF/pF), Ca^2+^ charge entry (pC/pF) during depolarization, and peak Na^+^ current (pA/pF) all normalized to cell size. N=96 and 73 cells, from 8 and 7 differentiations, and 47 cells from 11 donors. Box plots represent the median, interquartile range, and minimum/maximum values. Statistical significance was determined using Wilcoxon tests (ns: *p<0.05; **p<0.01, ***p<0.001 and ***p<0.001).

Despite low insulin secretion responses to glucose, both the SCβ D30 and SCβ D50 cells, identified by their expression of GFP under the *INS* promoter, exhibited larger exocytotic responses compared with primary β-cells, even after normalization to initial cell size (**Fig. 1E,F**). Sodium currents were also larger in both SCβ D30 and SCβ D50 than in primary β-cells (**Fig. 1F**, *third panel*). Similarly, Ca^2+^ influx was significantly higher in the SCβ D30 group than primary β-cells but did not differ between SCβ D50 and primary β-cells (**Fig. 1F**, *fourth panel*). These data indicate that SCβ-cells are not functionally mature, even though parts of the excitatory machinery and exocytotic processes appear highly active.

### Patch-seq links SCβ-cell dysfunction to altered ion channel transcripts and immature organelle-associated pathways

In a separate experiment (**Fig. 2A**), patch-seq analysis of SCβ-cell and primary β-cells confirmed the electrophysiologic differences observed above (**Fig. 2B**). We probed transcripts related to stimulus-secretion coupling and ion handling across β-cell populations (**Fig. 2C**). *CACNA2D1* (L-type Ca^2+^ channel auxiliary subunit) and *CACNA1E* (R-type Ca^2+^ channel) emerged as strong SCβ-enriched transcripts. Additional transcripts such as *ABCC8* (K_ATP_ channel subunit), *KCNH6* (voltage-gated K^+^ channel), *SCN9A* (voltage-gated Na^+^ channel), *CACNA1H* (T-type Ca^2+^ channel), *VDAC1* (MT Ca^2+^/ATP exchanger), and *SLC2A1* (GLUT1) were also among differentially expressed transcripts, although with somewhat lower confidence (**Suppl Table 1**). Canonical β-cell genes involved in insulin production, secretion, granule packaging, identity (e.g., *CHGB, MAFA, IAPP, PCSK2, INS*), and glucose metabolism (*ADCYAP1, G6PC2*) were enriched in primary β-cells. Conversely, *CHGA* and *HNF4A* exhibited higher expression in SCβ-cells (**Fig. 2D**; transcripts with high confidence are highlighted in bold red text, **Suppl Table 1**). Gene Set Enrichment Analysis (GSEA) comparing SCβ D30 and SCβ D50 transcripts revealed modest transcriptomic shifts over time in culture, particularly in pathways related to the mitochondrial respiratory chain complex and in ribosomal subunits (**Fig. 2E,F**, *top panels*).

**Figure 2:**
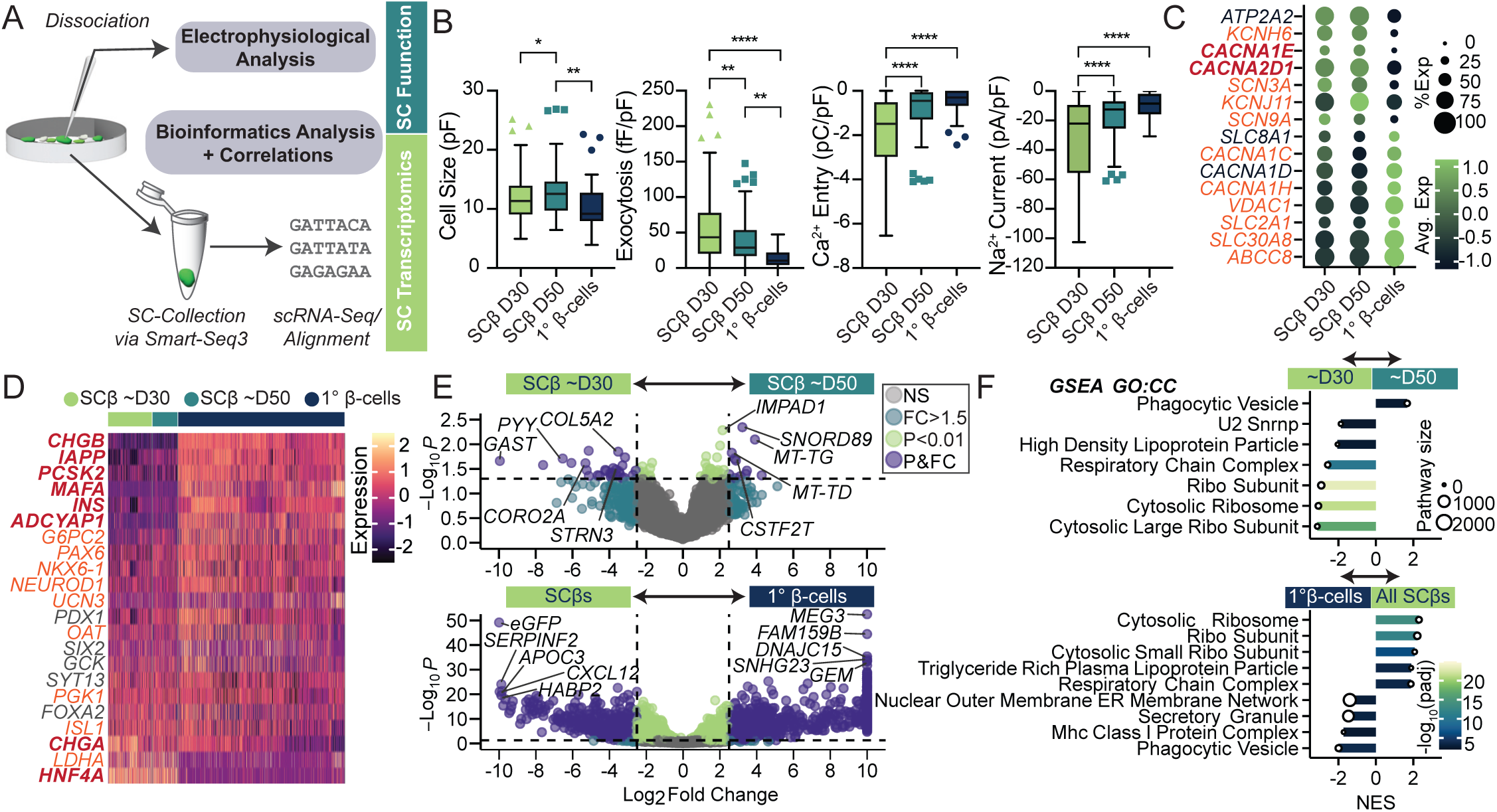
Patch-seq highlights functional and molecular differences between primary and SCβ-cells. (A) Schematic workflow illustrating single-cell RNA sequencing following electrophysiological and metabolic assessments. (B) Quantification of electrophysiological parameters across cell types. Statistical significance was determined using Wilcoxon tests (*p≤0.05; **p≤0.01; ***p≤0.001; ****p≤0.0001). N=159 and 110 cells, from 8 and 7 differentiations, and 60 cells from 10 donors. (C) Dot plot of key genes associated with β-cell ion channel activity. Dot size indicates average expression, and colour represents expression level. Transcripts shown in red indicate high-confidence differential expression, whereas orange indicates lower confidence. (D) Heatmap showing the canonical transcript expression profiles of functional and metabolic genes across SCβ D30, SCβ D50, and primary β-cell groups. Transcripts shown in red indicate high-confidence differential expression, whereas orange indicates lower confidence. (E) Volcano plot highlighting the top differentially expressed genes between SCβ D30 and SCβ D50 (*top*), and between all SCβ-cells and primary β-cells (*bottom*). Purple points indicate significantly enriched genes (log_2 fold change >2.5, FDR <0.05, or log_2 fold change >2.5, FDR <0.0001, respectively). (F) Gene set enrichment analysis (GSEA) results for Gene Ontology Cellular Components (GOCC), comparing SCβ D30 and SCβ D50 (*top*), and SCβ-cells and primary β-cells (*bottom*). Bars show normalized enrichment score (NES), colour encodes -log_10 (“adjusted “ p), and point size indicates pathway size.

As expected, large differences in transcriptomic profiles were observed between primary and SCβ-cells (**Fig. 2E,F**, *bottom panels*). GSEA revealed that phagocytic vesicle and secretory granule pathways were significantly enriched in primary β-cells, consistent with more functionally mature secretory phenotypes (**Fig. 2F**, *bottom panel*). Conversely, enrichment of pathways such as the respiratory chain complex is consistent with developing cell states.^27–29^ Additional analyses identified enrichment of cholesterol and lipid-related pathways (e.g. lipoprotein particle receptor binding, triglyceride-rich lipoprotein particle remodelling, and statin inhibition of cholesterol biosynthesis), which may be an additional feature of SCβ-cell immaturity (**Suppl Fig. S1**).

### Improved secretory granule and mitochondrial morphology with ageing of SCβ-cells does not equate with functional maturity

To investigate whether molecular signatures of immaturity were accompanied by structural and functional differences, we examined secretory granule and mitochondrial morphology in SCβ-cells at different stages of aging compared with primary β-cells. Transmission electron microscopy to assess ultra-structural features within SCβ D30 cells, D50 cells and primary islet-cells (**Fig. 3A**) showed that granule size was similar between groups but occupied a greater cytosolic area within SCβ D30 cells than in primary β-cells (**Fig. 3B**, *upper panels*). Granule density in the older SCβ D50 cells was lower than in the SCβ D30 cells, and somewhat lower than in primary β-cells (**Fig. 3B**, *upper panels*). While dense cores were less frequent, smaller, and reduced in solidity in the SCβ D30 group, these properties in the SCβ D50 group all closely resembled those in primary β-cells, consistent with the intermediate exocytotic phenotype observed above.

**Figure 3:**
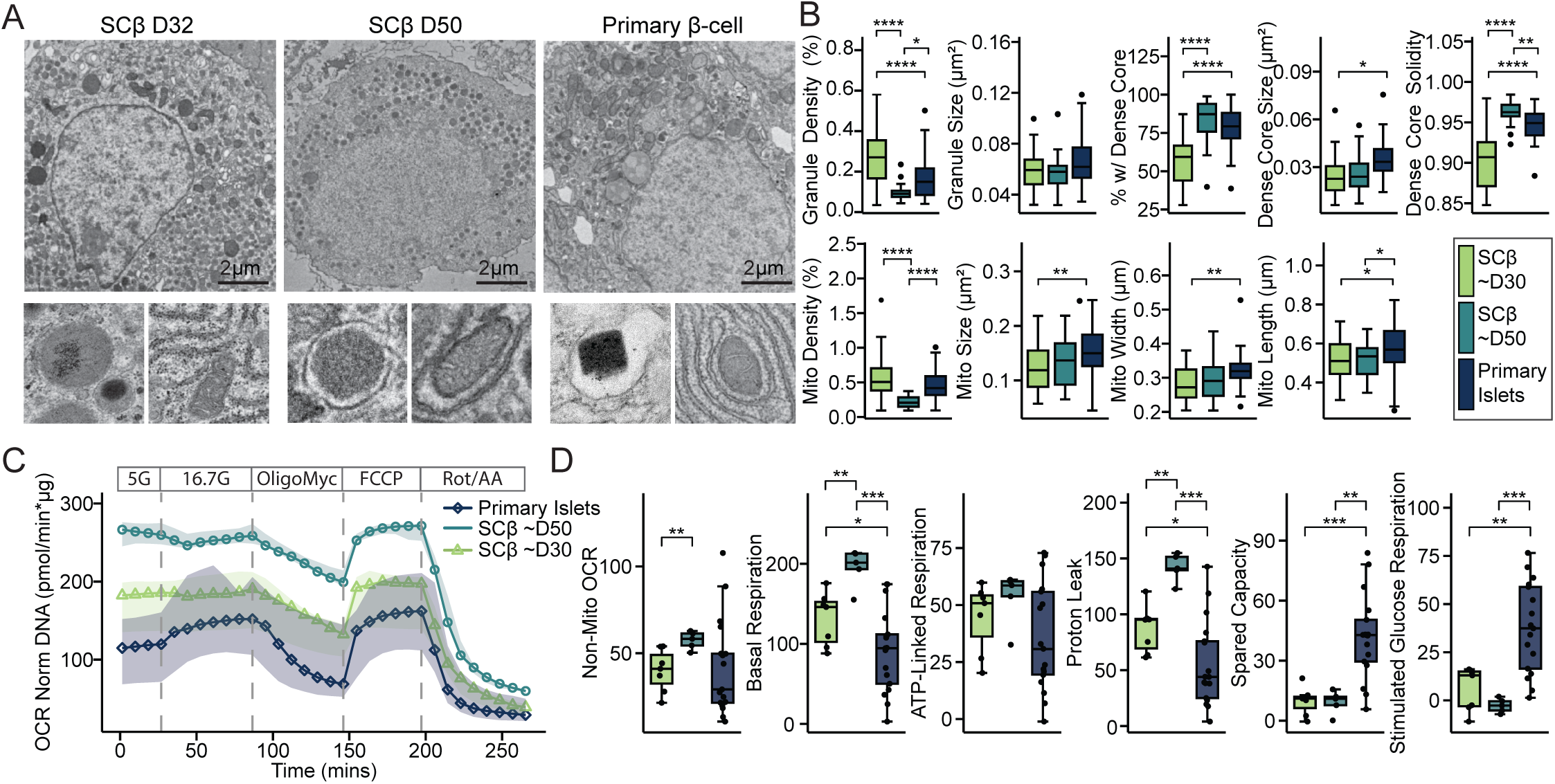
Morphological and functional assessments of SCβ-cell granules, mitochondria, and oxygen consumption rates. (A) Representative transmission electron microscopy images of SCβ D32, SCβ D50, and primary β-cells (scale bar, 2 μm). (B) Quantification of transmission electron microscopy images, including granule density (percentage of cytosol occupied by granules), granule size (μm^2^), percentage of granules with a dense core as determined by the Otsu method (%), dense-core size (μm^2^), and dense-core solidity (particle area divided by its convex hull area). Mitochondrial measurements were calculated similarly. Data are shown as median ± IQR (N=3 or 4 per group), with significance determined by Wilcoxon test (*p≤0.05; **p≤0.01; ***p≤0.001; ****p≤0.0001). (C) Oxygen consumption rates of SCβ D30 (N=8 differentiations), SCβ D50 (N=6 differentiations), and human islets (N=18 donors) in response to 5 mM glucose, 16.7 mM glucose, oligomycin A, FCCP, and antimycin A (3-4 technical replicates each). (D) Quantification of non-mitochondrial OCR, basal metabolism, ATP-linked respiration, proton leak, spare capacity, and glucose-stimulated oxygen consumption index from the data in panel C, with significance determined using Wilcoxon tests (*p≤0.05; **p≤0.01; ***p≤0.001; ****p≤0.0001).

Mitochondrial size, width, and length were smaller in SCβ D30 cells compared to primary β-cells, but no difference in mitochondrial morphology other than length was observed in the SCβ D50 group when compared to primary β-cells (**Fig. 3B**, *lower panels*). SCβ-cells had higher overall oxygen consumption rates than human primary islets (**Fig. 3C**). Both SCβ D30 and SCβ D50 cells exhibited elevated basal respiration compared with primary islets. Proton leak was also increased in both SCβ-cell groups relative to primary islets. SCβ D50 cells exhibited greater basal respiration and proton leak than SCβ D30 cells, indicating increased mitochondrial activity under resting conditions and reduced coupling efficiency. In addition, SCβ-cells had both lower spare respiratory capacity and glucose-stimulated oxygen consumption, indicating impaired glucose responsiveness. Thus, while the additional culture of the SCβ-cells resulted in secretory granule and mitochondrial morphologies more closely resembling those in primary β-cells, this alone was not sufficient for mature oxygen consumption or insulin secretory (**Fig. 1C,D**) phenotypes.

### Correlative analysis links key cellular programs with cell function

To increase the power of the study, we incorporated a dataset of previously published primary human islet patch-seq data (see Methods). We correlated each electrophysiological parameter with gene expression, highlighting transcripts with consistent, statistically significant associations across the SCβ-cells. We visualized the top 10 positively and negatively correlated transcripts, with cells ordered by increasing exocytosis, together with primary β-cells from the larger dataset for context (**Fig. 4A**). To contextualize these correlations, we performed GSEA using Gene Ontology (GO) Biological Process terms on genes ranked by their correlation with exocytosis within the SCβ-cells (**Fig. 4B**), identifying positive enrichment for early biosynthetic pathways (e.g. small ribosome subunit biogenesis, ATP synthesis coupled to electron synthesis, and cytoplasmic translation), and negative enrichment for pathways involved in secretion and stress handling (e.g. insulin processing, protein folding in ER, negative regulation of ER stress-induced apoptotic pathway) (**Fig. 4B**). This suggests that higher exocytosis is associated with more immature, biosynthetically active SCβ-cells, rather than a mature β-cell secretory state. Similar divergent transcript-electrophysiology correlations were seen across recorded parameters (**Suppl Fig. S2A**).

**Figure 4:**
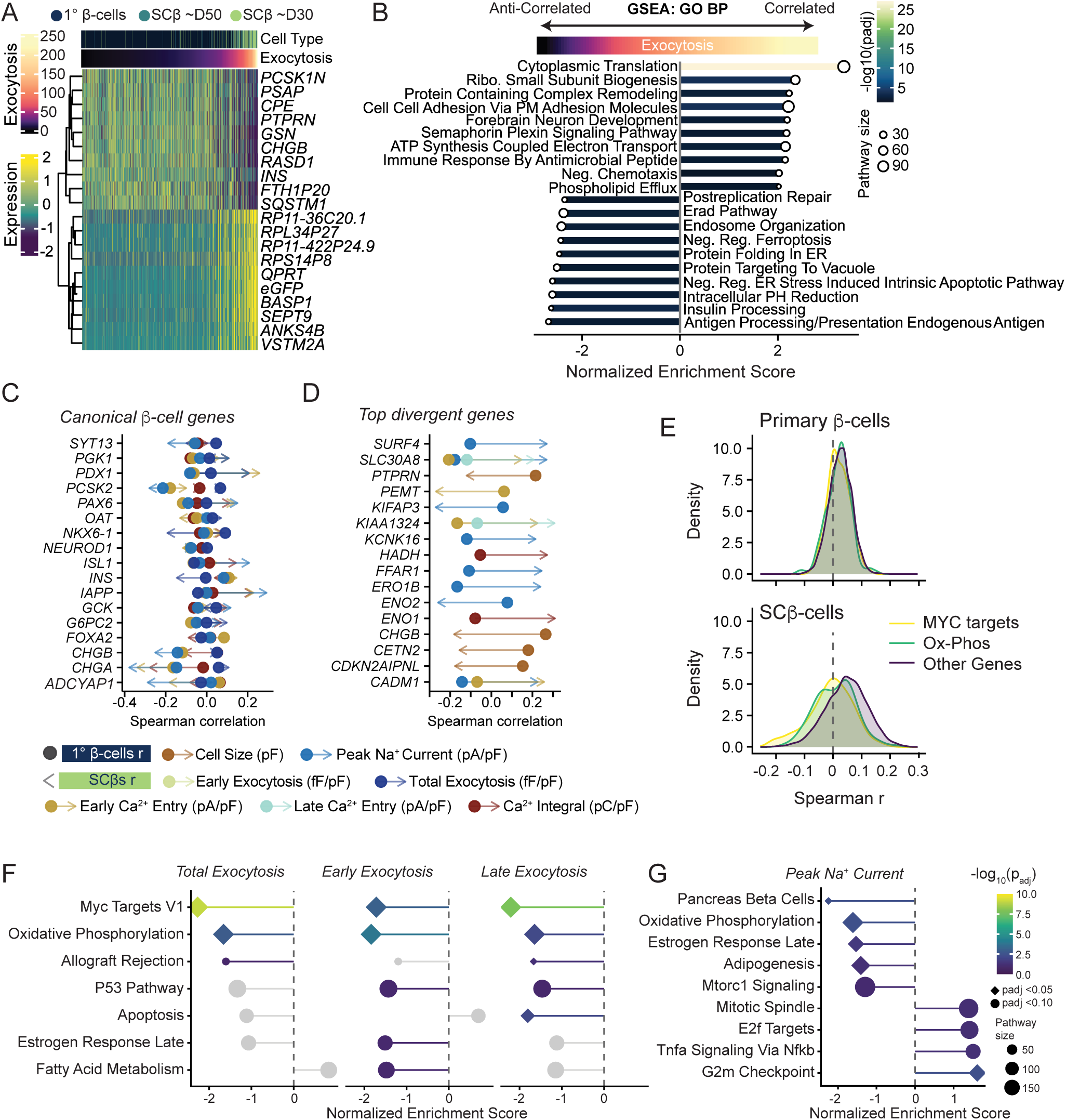
Transcript–electrophysiology correlation identifies shared and divergent gene programmes in primary and SCβ-cells. (A) Expression Heatmap (SCT-scaled Pearson residuals) of the top 10 positively and top 10 negatively correlated transcripts computed on all patched cells most associated with normalized exocytosis, filtered for significance (FDR<0.05), detection in >10% of cells, and a stable bootstrap range <0.10. SCβD30 (light green), SCβD50 (turquoise) and human islets (Navy) are ordered by increasing exocytosis. (B) GSEA (GOBP) of correlated and anticorrelated transcripts to normalized exocytosis in pooled samples of D30 SCβ, D50 SCβ and primary β-cells. Bars show normalized enrichment score, color encodes -log10(adjusted p) and point size indicates pathway size. (C) Transcript-electrophysiology relationships between primary and SCβ-cells, showing selected canonical β-cell genes. Arrow denotes spearman r for SCβ-cells, while circle denotes spearman r for primary β-cells. are shown. (D) Divergent associations transcript-electrophysiology relationships between SCβs and primary β-cells (ranked by |Δz|) prioritizing top transcript-electrophysiology relationships by limited heterogeneity between studies, boot strap stability and narrow confidence ranges. Associations were classified as divergent when |Δz| > 0.10, the confidence interval excluded 0, correlations showed either a sign flip or an absolute correlation of at least 0.15. Spearman correlations were calculated separately in SCβs and primary β-cells. Circle represents r_beta (meta-analytic estimate across five studies) while arrow represents r_SCβ. (D) Gene set enrichment analysis (GSEA) using Hallmark gene sets, ranked by Δz for total, early, and late exocytosis. All terms shown reach padj < 0.10; diamonds indicate padj < 0.05. (E) GSEA using Hallmark gene sets, ranked by Δz for for peak Na⁺ current. All terms shown reach padj < 0.10; diamonds indicate padj < 0.05.

To further identify divergent transcript-phenotype relationships that remain hidden when ranking genes based on within-population correlations alone, we computed Δz = z_SCβ − z_β using a random-effects meta-analytic framework combining primary β-cell estimates across five studies with multiple imputation to handle missing electrophysiology values (**Suppl Table 2**). While absolute correlations were modest within both populations (|r| ≤ 0.35), coherent patterns emerged. Some transcripts with conserved correlations between primary and SCβ-cells included *TNPO1*, *PBXIP1*, *ARHGAP12*, *MGST2*, *DDR1* (**Suppl Fig. S2B**). GSEA (GO Cellular Component) of these highlighted terms related to cell-cell adhesion, postsynaptic early endosomes, voltage-gated calcium channel complexes, and the Golgi lumen, among others (**Suppl Fig. S2C**). Several canonical β-cell and endocrine-specific transcripts, including *CHGA*, *CHGB*, *PDX1*, *IAPP*, and *ADCYAP1*, showed distinct correlations with electrophysiological parameters in SCβ-cells (arrowheads), relative to primary β-cells (circles) (**Fig. 4C,D**). Among the most divergent, *CHGB* showed a reversal for cell size, *ERO1B*, thought to promote protein folding in β-cells,^30^ and *CADM1*, which may regulate insulin secretion by mediating nerve–islet cell interactions^31^ showed strong divergence for Na^+^ current (**Fig. 4D**, **Suppl Table 2**).

GSEA on Δz-ranked genes (Hallmark gene sets) identified MYC targets and oxidative phosphorylation as a significantly negatively enriched for exocytosis (**Fig. 4E,F**), consistent with the impaired glucose-coupled oxidative metabolism observed in the SCβ-cells. Negative enrichment of MYC targets reflects reduced MYC target gene expression in the SCβ-cells with the highest exocytosis, consistent with evidence that c-Myc activity drives β-cell immaturity and impairs glucose-stimulated insulin secretion. Consistent with the GSEA finding, density plots of within-population correlations showed that both MYC target and oxidative phosphorylation genes are left-shifted relative to other genes in SCβ-cells (**Fig. 4E**), meaning higher expression of these genes associates with lower exocytosis, while no such shift is apparent in primary β-cells where correlations are compressed near zero.

For Na⁺ current, GSEA on Δz-ranked genes (Hallmark gene sets) identified Pancreas Beta Cells gene set as the most significantly enriched term (**Fig. 4G**). Canonical β-cell identity genes including *IAPP*, *PDX1*, *PCSK1*, and *ISL1* were negatively correlated with increased Na⁺ current activity in SCβ-cells, meaning SCβ-cells expressing higher levels of these mature identity markers had less Na^+^ channel hyperactivity (**Suppl Table 2**). Conversely, genes positively associated with higher Na⁺ current activity in SCβ-cells included *CHGA*, *STXBP1*, and *NKX2-2*, suggesting that a distinct transcriptional program tracks with the hyperactive electrophysiological state. Additionally, GSEA on Δz-ranked genes (GO Biological Process) indicated that Na^+^ current in primary β-cells was associated with regulation of ion concentration, oxidative phosphorylation, and endoplasmic reticulum protein folding and processing. Na^+^ currents in SCβ-cells were associated with pathways related to the maintenance of cell number, DNA biosynthetic processes, haematopoietic progenitor cell differentiation, sister chromatid cohesion and DNA-templated transcription initiation (**Suppl Fig. S2D**). Together, these findings indicate that SCβ-cells exist on a maturation continuum in which transcriptional and electrophysiological maturation are partially coupled, but the coupling is fundamentally reorganized relative to primary β-cells.

### Co-activity networks highlight fate-specific regulatory coordination

To assess regulatory programs in primary β-cells, SCβ-cells, and Stage 6, Day 1 stem cells (S6D1),^32^ we constructed SCENIC-based inferred co-activity networks (**Fig. 5A**, **Suppl Fig S3A**). While the primary β-cell network is modular, well-structured, and anchored by canonical regulators (PDX1(+), MAFA(+), NKX6-1(+), XBP1(+)), reflecting a mature transcriptional architecture, SCβ-cell networks were denser and less modular, suggesting a failure to establish coherent regulatory circuitry rather than the acquisition of an alternative mature state (**Fig. 5A**). S6D1 cells showed sparse, fragmented networks reflecting limited endocrine regulatory coordination (**Suppl Fig. S3A**). Nodes are colored by condition-specific influence score to visualize potential drivers within each regulatory state. Conversely, MNX1(+) (also known as HLXB9), a developmental transcription factor associated with early pancreatic specification,^33,34^ showed higher influence in SCβ-cells, further supporting the interpretation that SCβ-cells occupy a transcriptionally immature or progenitor-like regulatory state.

**Figure 5:**
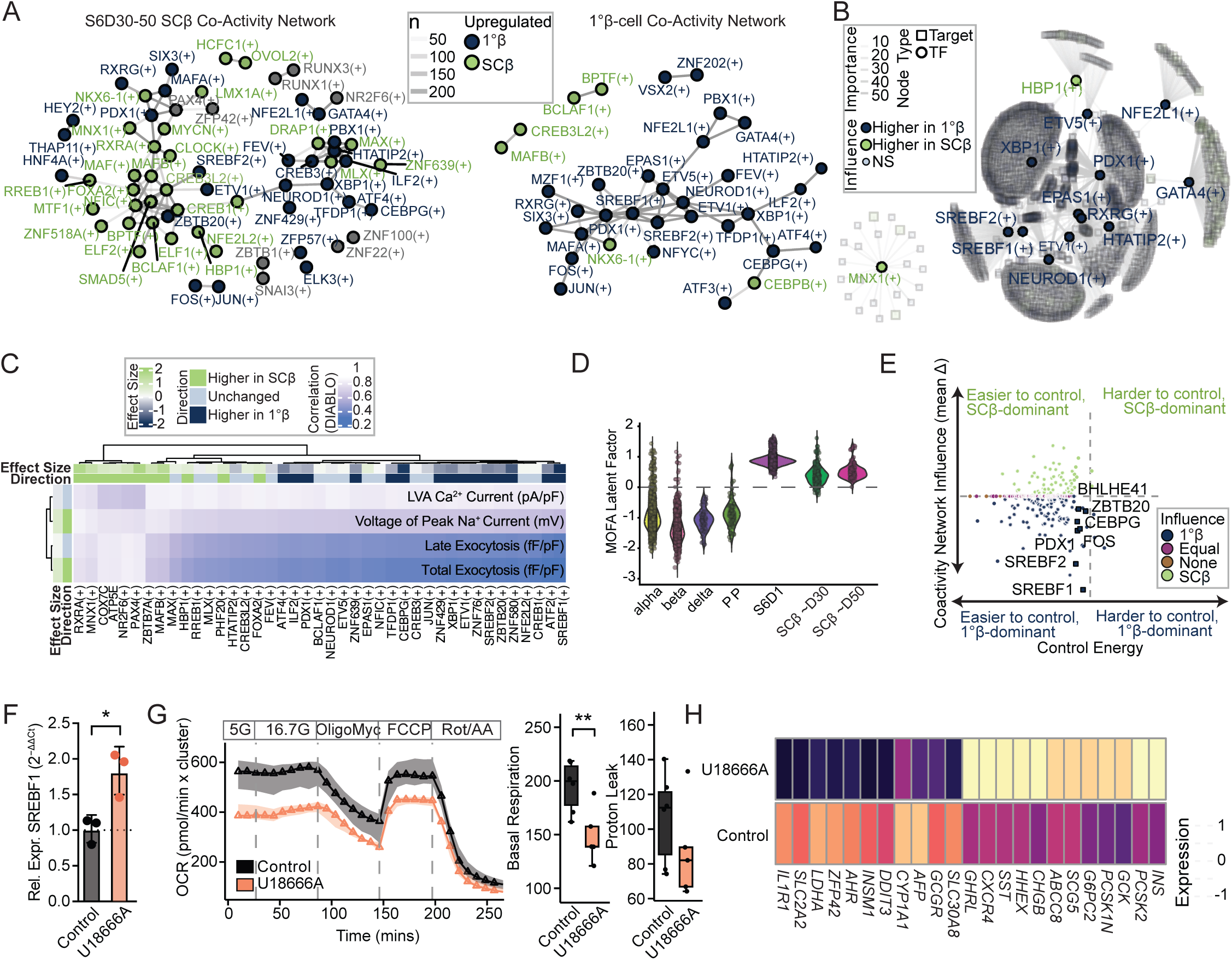
SCENIC co-activity and multi-omics integration reveal targetable SCβ-cells regulatory disorganization. (A) Bootstrapped co-activity networks derived from SCENIC AUC values for SCβ cells (*left*), and primary β-cells (*right*). Edges represent reproducible regulon co-activity across bootstrap iterations (present in ≥50% of iterations). Nodes are colored according to condition-specific influence. (B) Top 10 transcription factors (TF) ranked by bootstrapped Δ Influence between SCβ-cells and primary β-cells, together with their top target genes. Nodes are shaped by type (TF or target) and colored by direction of differential expression. Transcription factors with greater influence in SCβ-cells are broadly rewired and often regulate targets that are also upregulated in SCβ-cells, suggesting active roles in altered gene regulatory programs. © Cross-block correlation heatmap from consensus DIABLO supervised multi-omics integration. Rows show electrophysiology parameters and columns show SCENIC regulon AUC features and mRNA features selected by DIABLO (present in ≥80% of 50 imputation models with ≥90% sign consistency). Cell color encodes the mean cross-block DIABLO correlation between each electrophysiology–transcriptomic feature pair, averaged across valid imputation models. Row and column annotation bars show standardised effect size (estimate/SD of outcome, equivalent to Cohen’s d) and direction from linear mixed models (SCβ- vs. primary β-cells, Differentiation as random effect), with a shared diverging color scale across both annotation tracks. Features with padj > 0.05 are labeled ‘Unchanged.’ Hierarchical clustering uses Euclidean distance with Ward.D2 linkage. Current amplitude parameters were sign-flipped prior to analysis so that more positive values indicate greater electrophysiological activity. (D) Consensus MOFA2 latent factor scores by cell type and differentiation stage. Scores represent the mean across 50 imputation runs of the primary separation factor (the factor with maximum standardized effect size separating SCβ-cells and S6D1 cells from primary endocrine cells) after sign alignment across runs. Each point represents a single cell; violin shapes show the distribution. Dashed line indicates zero. (E) Co-activity network influence versus control energy for candidate transcription factors. Each point represents a transcription factor, plotted by control energy (x-axis) and bootstrapped mean influence difference (change in bootstrapped PageRank influence) between primary and SCβ-cell co-activity networks (y-axis). Labelled transcription factors were selected based on high composite scores that integrate influence stability, differential AUC activity, and control energy. Colour indicates the cell type in which the transcription factor is more influential, and shape denotes selected candidates. (F) *SREBF1* transcript expression measured by qPCR in SCβ-cells treated with U18666A versus control, normalized to the housekeeping gene PPIA (Student’s t-test, P=0.0149, N=3 and N =3, respectively). (G) Oxygen consumption rates of SCβ-cells treated with U18666A (N=3) or control (N=3) in response to 5 mM glucose, 16.7 mM glucose, oligomycin A, FCCP, and antimycin A (3–4 technical replicates each). (H) Treatment-responsive genes identified by NanoString profiling and limma analysis, with batch included in the design matrix and Benjamini–Hochberg multiple-testing correction applied (adjusted P<0.05; N=6).

To investigate whether specific transcription factors differentially coordinate gene regulatory activity between primary and SCβ-cells, we applied an influence propagation analysis which identified transcription factors with divergent network influence, and visualized their downstream gene targets (**Fig. 5B**, **Suppl Fig. S3B**, **Suppl Table 3**). Notably, SREBF1(+) and SREBF2(+), regulators of lipid and cholesterol homeostasis, showed higher network influence in primary β-cells (**Fig. 5B**) and co-activated within a broader β-cell-enriched regulatory module that includes PDX1(+), NEUROD1(+), MAFA(+) and NKX6-1(+) (**Fig. 5A**). Consistent with this, the target genes of these β-cell-dominant transcription factors, including genes involved in insulin processing, secretory granule maturation, and glucose metabolism, were also differentially expressed at the transcript level, with higher expression in primary β-cells, providing evidence across independent analyses that the loss of regulatory influence in SCβ-cells is accompanied by a coordinated transcriptional deficit in the downstream programs these transcription factors normally coordinate. This pattern supports the idea that SREBF1(+) is part of a coherent co-activated β-enriched regulatory module, which also includes NEUROD1(+), MAFA(+) and NKX6-1(+) (**Fig. 5A**, *right panel*). The reduced network influence of these canonical β-cell regulators in SCβ-cells indicates that the failure of SCβ-cells to achieve mature function reflects not just the absence of individual regulators but a broader loss of higher-order regulatory integration that characterizes functional β-cell identity. We also observed that this core group of regulons coordinates organized network co-activity by differentiation/maturation, mitochondrial/metabolic, vesicle/transport, and neuroendocrine gene programs (**Suppl Fig. S3B**).

### Integrated analysis identifies dysregulated SCβ-cell transcript-electrophysiology coupling

To understand how transcriptomic and electrophysiological features co-vary and discriminate between SCβ-cells and mature endocrine cells, pseudobulked data aggregated by differentiation batch or human islet donor and cell type were analyzed using DIABLO, using the expanded patch-seq dataset (see Methods). Hierarchical clustering of cross-block correlations with electrophysiological properties revealed two broad patterns. Most transcription factor regulons, including SREBF1(+), SREBF2(+), NEUROD1(+), and PDX1(+)) were more active in β-cells and showed weak positive correlations with exocytotic capacity (normalized total and late exocytosis) and strong positive correlations with Ca^2+^ and Na^+^ channel activities (**Fig. 5C**). In contrast, a smaller cluster of regulons, including PAX4(+), RXRA(+), and MNX1(+) were more active in SCβ-cells and strongly correlated across exocytotic parameters. The latter (MNX1(+)), also identified as having higher network influence in SCβ-cells by co-activity analysis (**Fig. 5A,B**), represents convergent evidence across independent analytical frameworks. MAFB(+), also more active in SCβ-cells, formed a separate cluster showing strong correlations with low voltage-activated (LVA) Ca²⁺ current and voltage of peak Na⁺ current, electrophysiological features that also correlate with the primary β-cell dominant transcription factor module, suggesting that MAFB may represent a partially matured regulatory program in the SCβ-cells.^35^ Normalized LVA Ca^2+^ current and voltage of peak Na^+^ current did not differ significantly between SCβ-cells and primary β-cells despite loading strongly on the discriminant component, suggesting that while these electrophysiological properties co-vary with transcriptional state across the dataset, they may not represent a primary axis of immaturity between these two populations specifically (**Fig. 5D**). Together, these findings indicate that primary β-cells and SCβ-cells differ not only in the magnitude of transcriptional and electrophysiological activity, but in how those layers are coupled: mature β-cell identity transcription factors associate with ion channel properties, while SCβ-cell dominant regulatory programs associate with exocytotic overactivity.

To define sources of hidden but shared variation across cell states, we applied unsupervised analysis using MOFA2, performed at the single-cell level. This identified a latent factor structure that separated SCβ-cells from primary endocrine cells, despite providing no supervision or cell type labels during factor learning, reinforcing that integrated regulatory, transcriptional and electrophysiological features distinguish these populations (**Fig. 5D**). Primary α, β, δ and PP cells form clusters in UMAP space, while SCβ-cells form a distinct group, (**Suppl Fig. S3C**) suggesting a transcriptionally coherent but divergent identity. S6D1 cells, representing an earlier differentiation state, do not overlap with D30 or D50 SCβ-cells, suggesting that SCβ-cells represent a biologically distinct differentiation state relative to earlier protocols. This embedding reinforces that MOFA2 factors reflect underlying cell identity and that SCβ-cells, while distinct from β-cells, do not simply recapitulate a less mature progenitor state but represent a biologically distinct identity. The primary latent factor separated mature endocrine cells from all SCβ populations, with the highest-loading features including canonical β-cell identity regulons and electrophysiological parameters on the mature side (**Suppl Fig. 3D,E**, **Suppl Table 4**).

### Network control analysis prioritizes drivers of SCβ-cell maturation

To identify candidate regulators of SCβ-cell maturation, we ranked transcription factors by their predicted ability to drive SCβ-cells toward a primary β-cell phenotype. Control energy was calculated by modelling transcription factor-target interactions as a linear dynamical system and computing the energy required to shift multiomic patterns towards a primary β-cell state, via forced response simulation, following principles from control theory (**Fig. 5E**, **Suppl Table 5**). Transcription factor rankings were stabilized using 100 bootstrap resamplings of cells. We characterized each factor by its regulatory drive (predicted control energy) and its differential impact between cell types (mean Δ influence, which was higher in primary *versus* SCβ-cells). Consistent with previously computed networks, SREBF1, SREBF2, and PDX1 were top candidates, exhibiting both high β-cell-dominant network influence and relatively low predicted control energy to shift SCβ-cells towards a more mature state.

Emerging as a top-ranked maturation candidate with high network influence and low predicted input energy was SREBP1 (sterol regulatory element-binding protein-1, encoded by the *SREBF1* gene), an ER-tethered transcription factor regulating cholesterol and lipid homeostasis. We tested this axis by perturbing cholesterol pathways by treating differentiating SCβ-cells from D15 to D30 of differentiation with 1 µM U18666A, a drug shown to increase *SREBF1* expression and subsequent activity.^36^ U18666A-treated SCβ-cells had increased SREBP1 expression (**Fig. 5F**) and exhibited a lower-shifted oxygen consumption profile with lowered basal respiration (**Fig. 5G**). Transcriptional profiling showed significant increases in *INS*, *CHGB*, *GCK* and *PCSK2* expression, with simultaneous decreases in *LDHA* (a marker of glycolytic immaturity in β-cells), *DDIT3* (unfolded protein response), and *ZFP42* (marker for undifferentiated stem cells) (**Fig. 5H**, **Suppl Table 6**). These data validate the SREBP1 axis as a functional maturation target, while also suggesting that full maturation may require broader targeting of TFs that rewire whole regulatory networks, rather than targeting individual marker genes.

## DISCUSSION

SCβ-cell immaturity is not simply a deficit in individual identity markers, but a failure to establish the higher-order regulatory integration that couples transcriptional programs to coordinated electrophysiological and metabolic function. We interrogated functional, molecular and regulatory phenotypes of SCβ-cells, systematically integrating these aspects of cellular function to reveal a holistic understanding of their immaturity. This approach is suitable for smaller, deeply annotated datasets, where cross-modality integration reduces the limitations of individual measurements. While electrophysiological features like exocytosis and cell size differ between SCβs and primary β-cells, significant within-population variability shows no single feature defines either cell type. Consistent findings from both unsupervised (MOFA2) and supervised (DIABLO) methods support SCβ-cell separation from primary cells and the role for SREBP axis dysregulation.

This study, like others, has consistently reported low glucose-stimulated insulin release in SCβ-cells, which has been attributed to glycolytic and anaplerotic gaps necessary for coordination of excitability.^6,11,14,26^ While our SCβ-cells show incremental transcriptional and functional maturation with increasing differentiation stage and time in culture, these remain distinct from primary β-cells across all measured modalities. The larger Na^+^ and Ca^2+^ currents and exocytotic responses we find in SCβ-cells indicate that, although the ionic machinery necessary for stimulus-secretion coupling is present and perhaps overactive, cellular coordination is lacking. In the SCβ-cells, exocytosis tracked positively with biosynthetic pathways, and was negatively associated with MYC target, oxidative phosphorylation gene programs, mature secretory and ER-folding pathways. In primary β-cells, these programs showed no relationship with exocytosis, consistent with a more uniform expression of pathway transcripts across the mature population. This suggests that within the SCβ-cell population, the most dysregulated cells are also the most metabolically immature.

Supporting the hypothesis that β-cell maturity not only requires correct transcript expression but also coordinated network activity, many canonical identity factors (e.g. *PDX1, PAX6, GCK, SIX2, SYT13, FOXA2*) were not differentially expressed between primary and SCβ-cells, but primary β-cells exhibited a more structured network centred around canonical maturity factors like PDX1(+), MAFA(+), SREBF1(+), NEUROD1(+) and NKX6-1(+). Within a denser, and less modular, network defining SCβ-cells, MNX1(+), a transcription factor required for early pancreatic specification and whose mutations cause neonatal diabetes in humans,^33,34^ showed consistently higher network influence in SCβ-cells relative to primary β-cells. This was independently confirmed by DIABLO multi-omics integration, where it emerged as part of a cluster of SCβ-cell-dominant regulons correlated with exocytotic overactivity, and suggests retention of an early pancreatic progenitor-like regulatory state. The co-occurrence of this with the PAX4 regulon in the SCβ-cell-dominant DIABLO cluster is notable here since PAX4 mis-expression in mature β-cells increases c-Myc and promotes dedifferentiation.^37^ Thus, despite transcriptional similarities, SCβ-cells retain a partially resolved developmental regulatory program, in which progenitor-associated transcription factors remain active and prevent a mature regulation of the secretory machinery that characterizes primary β-cells.

Transcript-electrophysiological relationships were fundamentally different between SCβ-cells and primary β-cells. While primary β-cell correlates were enriched in oxidative phosphorylation, calcium regulation, and ER protein-handling pathways, SCβ-cell correlates were more closely associated with developmental and proliferative programs. This suggests that similar functional readouts are embedded in different biological contexts in the two cell types. Mitochondrial function assays further reinforced this functional immaturity, with in vitro-matured cells showing an increased basal oxygen consumption rate compared with their nascent counterparts. The absence of glucose-stimulated respiration and elevated proton leak suggests mitochondrial uncoupling rather than mature oxidative capacity. This paradox of more mitochondrial activity without coordinated glucose-responsive secretion echoes the transcript-electrophysiology finding that metabolic programs are not appropriately coupled to secretory function in SCβ-cells, as suggested by others^6^, and suggests that extended culture time alone is insufficient to resolve the underlying regulatory decoupling. This could be due to culture conditions that are insufficient to support the transition from an immature regulatory network to a more coordinated one. The enrichment of lipoprotein and cholesterol-related pathways in SCβ-cells, together with subsequent prioritization of *SREBF1/SREBF2* and the positive impact of experimental perturbation of cholesterol handling, suggest that sterol metabolism and lipid regulation may be mechanistically linked to the immature β-cell state and perhaps constitute an underappreciated axis contributing to this immaturity.

While the mean values of electrophysiological features such as exocytosis and cell size differ significantly between SCβ-cells and primary β-cells, the variability in their distribution highlights the limitations of using a single measurement to define the cell types. Integration of these different data layers confirmed that our SCβ-cells are distinct from primary β, α, δ and PP cells, with SCβ-cells exhibiting features of multiple cell states and persistent progenitor-like features across transcriptomic, regulatory, and functional layers. To produce biologically meaningful hits to guide differentiation targets, control energy modelling identified SREBF1(+), SREBF2(+), PDX1(+), FOS(+), CEBPG(+), ZBTB20(+), and BHLHE41(+) as top candidate regulators capable of efficiently rewiring SCβ-cells. Although PDX1 was identified in this analysis and is a hallmark of β-cell identity, its high control energy suggests that reprogramming via PDX1 targeting alone may be inefficient and difficult. Indeed, others have shown that Pdx1 is insufficient for β-cell reprogramming, requiring co-expression of Ngn3 and MafA to achieve the full transition *in vivo*.^38^ This highlights the value of prioritizing regulators with favourable network properties over canonical identity markers alone, and we chose to test this by asking whether activating the *SREBF1*/sterol/lipid axis could shift SCβ-cells toward a more mature β-cell state, as this axis lies at the intersection of β-cell identity and lipid metabolism.^39^ Utilizing U18666A, which depletes ER-cholesterol,^36^ produced a pro-maturation phenotype, increasing *SREBF1* transcript, signifying pathway engagement. OCR/basal respiration was decreased in the U18666A-treated cells. Despite a modest increase in *SREBF1* transcript, and in agreement with SREBP1’s high network centrality, we saw widespread effects in β-cell specific transcripts related to mitochondria (*PCSK2, GCK, PCSK1N, G6PC2, SCG5, ABCC8, HHEX*) and β-cell identity (*INS, CHGB*).

Taken together, these results suggest that SCβ-cell immaturity arises from the failure of established functional regulatory network hierarchies that integrate electrophysiology, metabolism, secretion, and maturation. This work provides a blueprint for refining SCβ-cell transcriptional landscapes, including a ranked list of potential targets, and supports the broader conclusion that targeting network-level regulators may be more effective than focusing on single maturity markers alone. The integrated multi-omics framework presented here provides an approach that could be adapted to evaluate and improve other differentiation protocols. Our prioritized list of targets that map molecular drivers to functional phenotypes could help push SCβ-cells even closer to their primary β-cell counterparts, although this would need to be tested across multiple differentiation protocols.

### Limitations and Future Directions

Our findings aim to provide a comprehensive picture of the immature SCβ-cell phenotype; however, several limitations in technology still exist. While Patch-seq provides powerful single-cell resolution, low throughput makes it difficult to overcome the noise inherent to scRNA-sequencing technology. To overcome this limitation, we performed correlative analysis from both cells collected in this study, and integrated cells from previous studies to get a broad sense of cellular identities found to be conserved despite different sequencing strategies used. However, combining multiple cell sources still led to confounding batch effects. By utilizing a multi-modal approach with both supervised and unsupervised methods, we confirmed that the conclusions were robust and similar across analyses. Integration with epigenomic or proteomic data could further elucidate the post-transcriptional regulation contributing to the observed functional decoupling. The network control analysis used here approximates transcription factor input energy via a forced-response simulation of a linear dynamical system, rather than computing the true minimum control energy via the controllability Gramian. This approach identifies relative rankings of candidate regulators but does not provide theoretically optimal energy estimates and assumes a linear approximation of gene regulatory network dynamics that may not fully capture the nonlinear nature of transcriptional regulation. The nominated targets should therefore be interpreted as hypothesis-generating rather than definitive predictions, and experimental validation through CRISPR, inducible overexpression systems, or small-molecule experiments will be necessary to determine their true reprogramming potential in SCβ-cells.

## MATERIALS AND METHODS

### Cell culture

Human islets were isolated at the Alberta Diabetes Institute (ADI) IsletCore (www.isletcore.ca) as described.^40^ Islets were hand-picked and dissociated to single cells using enzyme-free Hanks’-based Cell Dissociation Buffer (Thermo Fisher Scientific, Cat#13150-016) and cultured in a glucose concentration of 5.5 mmol/L, DMEM with L-glutamine, 110 mg/L sodium pyruvate, 10% FBS, and 100 U/mL penicillin/ streptomycin at 37°C for 1-3 days. Donors gave informed written consent for research, and all studies were approved by the Human Research Ethics Board at the University of Alberta (Pro00013094; Pro00001754). SCβ-cells were generated as outlined^24^ and cultured for an additional 23–35 days (designated as ‘D30 SCβ-cells’) or for 45–54 days (designated as ‘D50 SCβ-cells’). Experiments presented in **Fig 5F-H** included 9µM WIKI4 (MedChemExpress; Cat# HY-16910) during stage 4, days 7-9.^41^ SCβ-cells were dissociated as described above with the addition of 10 µM ROCKi (Y-27632, cat # 72302) in the media and plated on a tissue culture-treated 35 mm dish and kept for 1-3 days.

### Glucose-stimulated insulin secretion assays

Dynamic glucose-stimulated insulin secretions were done after at least one overnight culture of human islets or SCβ-cell clusters using the Biorep Perifusion Machine (ver peri4.2) machine. Krebs-Ringer bicarbonate HEPES (KRBH) was prepared freshly on the day of the experiment containing 115 mM NaCl, 5 mM KCl, 2.5 mM CaCl₂, 1 mM MgCl₂, 10 mM HEPES, 24 mM NaHCO_3_, and 0.1% bovine serum albumin (BSA) and warmed to 37°C and pH was balanced to 7.4 with NaOH. Before sample collection, the perifusion apparatus was primed with a flow rate of 100 µL/min for the remainder of the protocol. 45 washed Islets or clusters were loaded into the chambers. The protocol consisted of a 30-minute pre-incubation step, followed by 5 minutes with 2.8 mM glucose KRBH, 28 minutes of 16.7 mM glucose KRBH, 18 minutes at 2.8 mM glucose KRBH and finally 8 minutes with 20 mM KCl in KRBH. Total insulin content was determined by running chambers dry after completion of the protocol, the filter paper containing islets or clusters was then transferred to a 1.5 mL Eppendorf tube contain 500 µL of Acid ethanol (150 mL 95% ethanol, 47 mL acetic acid and 3 mL concentrated HCL).

For static insulin secretion experiments,^42^ 15 islets/clusters were washed and preincubated (2.8 mM KRBH for 1 hr at 37 °C, 5% CO_2_) and placed into Eppendorf tubes (each group containing 3 technical replicates). 500µL of 2.8 mM KRBH was added to the tubes for 1 hr at 37 °C, 5% CO_2_ with lids open, followed by gentle inversion and centrifugation at 1000 rpm. The supernatant was then collected. 500 µL of 16.7 mM KRBH was added to the tubes next, for 1 hr at 37 °C, and supernatant collection was repeated. Finally, 500 µL of acid ethanol was added for collection of insulin content. Samples stored at −20°C were then assayed for Insulin using Alpco Stellux human insulin ELISA.

### Transmission electron microscopy

For ultrastructure analysis, cells from primary human islets (3 donors), D30 and D50 SCβ-cells (3 differentiations each) were dissociated and fixed in a buffer of sodium Cacodylate Buffer (0.1M, pH 7.4, from Electron Microscopy Sciences, cat#11650), 2.5% Glutaraldehyde (Electron Microscopy Sciences, cat#16350), 2% Paraformaldehyde (16% aqueous solution, Electron Microscopy Sciences, cat#15700), and 2mM Calcium Chloride (CaCl_2_). Cells were then pelleted, resuspended, and fixed at 37℃ for 20 mins and room temperature for 40 mins, followed by 24 hours at 4℃ to ensure the ultrastructure was maintained. After initial fixation, the cells were washed in sodium 0.1M Cacodylate Buffer (NaCB) for 10-15 mins (3 times) followed by post-fixing in freshly made 1% K_4_FeCN_6_ and 1% OsO_4_ in 0.1M NaCB for 1 hour on ice. The cells were again washed in 0.1M NaCB (overnight), washed with 0.1M Na Acetate (3 mins x 3 times), stained en bloc with 2% uranyl acetate (1hr RT) and washed with 0.1M Na Acetate (5mins X 2). This block was dehydrated in an ethanol concentration series to remove water, then embedded in Spurr’s resin for ultrathin sectioning. The blocks were then sectioned to 60 to 90 nm thickness with a diamond knife (45° angle) on an ultramicrotome and contrasted with uranyl acetate and lead citrate, and carbon coated on a 300-mesh copper grid. The stained sections were imaged using JEOL 2100, focusing on granules, ER, and mitochondria.

### Oxygen consumption measurements

Mitochondrial function was assessed using an Agilent Seahorse XFe24 Analyzer with islet capture microplates (Agilent). On the day of the assay, 50 SCβ-cell clusters or human islets were transferred onto XF islet capture plates in assay medium (Agilent DMEM supplemented with 1% heat-inactivated FBS, 2.8 mM glucose, 2 mM pyruvate, and 2 mM glutamine) and secured with islet capture screens to prevent movement. Plates were then incubated at 37^∘^C in a non-CO_2_ incubator for 1 h. The Mito Stress Test protocol was then performed by sequential injection of glucose (basal condition: 5 mM; stimulatory condition: 16.7 mM), oligomycin (5 μM), FCCP (3 μM), and rotenone/antimycin A (5 μM each). Oxygen consumption rate (OCR) was measured at baseline and after each injection using Wave software. Clusters and islets were then collected for normalization to DNA content.

### Gene expression by qPCR and NanoString

RNA extraction and reverse transcription quantitative PCR (RT-qPCR) were performed as described previously, and gene expression was normalized to the housekeeping gene PPIA (Thermo Fisher, Cat. 4331182, assay ID: Hs04194521_s1). SREBF1 expression was measured using a TaqMan assay (Thermo Fisher, Cat. 4448892, assay ID: Hs02561944_s1). For Nanostring assays, samples were analyzed on a NanoString nCounter SPRINT Profiler and processed using nSolver 4.0 for relative gene expression. Data were normalized to six housekeeping genes: *B2M*, *GAPDH*, *GUSB*, *HPRT1*, *POLR2A*, and *TBP*.

Differential expression analysis of log_2-transformed NanoString counts was performed using the limma package in R. A linear model including group and batch terms was fitted using the design formula ∼ 0 + group + batch, and contrasts comparing U18666A, and control were tested with empirical Bayes moderation (eBayes(trend = TRUE)). Multiple testing correction was performed using the Benjamini-Hochberg false discovery rate (FDR) method, and genes with adjusted p<0.05 were considered significant. Downstream visualization included heatmaps and volcano plots.

### Electrophysiology

For whole-cell patch-clamp recordings, fire-polished thin-wall borosilicate pipettes coated with Sylgard (3–5 MΩ) were used. Pipettes were filled with intracellular solution containing (in mM): 125 Cs-glutamate, 10 CsCl_2_, 10 NaCl, 1 MgCl_2_⋅6H_2_O, 0.05 EGTA, 5 HEPES, 0.1 cAMP, and 3 MgATP (pH 7.15 adjusted with CsOH). On the day of the experiment, media were replaced with the appropriate bath solution containing (in mM): 118 NaCl, 20 TEA, 5.6 KCl, 1.2 MgCl_2_⋅6H_2_O, 2.6 CaCl_2_, 5 HEPES, and either 1, 5, or 16.7 mM glucose (pH 7.4 adjusted with NaOH) in a heated chamber maintained at 32–35°C.

Electrophysiological measurements were collected using a HEKA EPC10 amplifier and PatchMaster software (HEKA Instruments, Lambrecht/Pfalz, Germany) within 5 min of break-in. Measured parameters included cell size (pF) as total membrane capacitance before the depolarization protocol, and subsequently used as a normalization control; early, late, and total exocytosis as capacitance increases (fF/pF) in response to a series of ten 500 ms depolarizations from −70 to 0 mV; and Na^+^ and Ca^2+^ currents elicited by a single 500 ms depolarization from −70 to 0 mV, collected as peak Na^+^ current (pA/pF), and early and late Ca^2+^ currents (pA/pF) and Ca^2+^ charge entry (pC/pF). Low- and high-voltage activated Ca^2+^ currents (LVA, HVA; pA/pF) were determined from voltage ramps. Quality control criteria included seal resistance >10 GΩ and access resistance <20 MΩ. Data were analyzed using FitMaster (HEKA Instruments) and Prism 10.0 (GraphPad Software, San Diego, CA).

To collect cells for scRNAseq, immediately following recording the patch pipette was withdrawn and replaced with a wide-bore collection pipette (0.2–0.5 MΩ) containing 0.5 μL lysis buffer. Cells were collected by gentle suction with visual confirmation and transferred into 8-strip PCR tubes containing 3 μL lysis buffer on ice, then stored at −80^∘^C until cDNA and library preparation.

### Single-cell RNA sequencing

Following patch-clamp recording, cells were collected into RNase-free 0.2 mL strip tubes and assembled into 96-well plates (Bio-Rad, RC9601 and MSA5001). cDNA libraries were generated from collected cells using a modified version of the Smart-seq3 protocol, as previously described.^43^ The mRNA was primed with an anchored oligo-dT primer (Smartseq3_OligodT30VN) and reverse transcribed using the Smartseq3_N8_TSO template-switching oligo, followed by PCR amplification (24 cycles). Libraries were then generated from amplified cDNA by tagmentation with Tn5. Libraries were sequenced on a NextSeq 500 platform (Illumina) using paired-end 75 bp reads to an average depth of 1 million reads per cell.

Single-cell RNA sequencing data were aligned to the GRCh38 reference genome using STAR^44^ and processed using the Seurat R package (version 4.0; https://satijalab.org/seurat/). Putative doublets were systematically identified using Scrublet and excluded from downstream analysis. Ambient RNA contamination was removed using the DecontX package prior to downstream analysis. Quality control metrics included mitochondrial gene content, ribosomal protein gene content, number of detected genes (nFeature_RNA), and total transcript counts (nCount_RNA). Cells with fewer than 200 detected genes were considered low quality and removed. Cells with a high proportion of mitochondrial transcripts (>40%) were also excluded.

### Bioinformatic methods

#### Integrating Patch-seq Datasets

Patch-seq scRNA-seq data from the current study and five previously published primary islet patch-seq datasets (obtained from the NCBI Gene Expression Omnibus (GEO) and Sequence Read Archive (SRA) under accession numbers GSE124742, GSE164875 and GSE270484; at PancDB (https://hpap.pmacs.upenn.edu/); and at Human Cell Atlas Data Explorer under project label CryoPancreaticIsletCellPatchSeq) were integrated into a single Seurat object. For each study separately, ambient RNA contamination was removed using decontX (celda package), replacing raw counts with decontaminated count estimates. Putative doublets were identified and removed using Scrublet (expected doublet rate = 0.06; cells with predicted_doublets = TRUE excluded), with cells exceeding 50% ambient contamination also removed. All downstream analyses used decontaminated counts as the RNA assay input. The S6D1 dataset (obstained from GSE120522) underwent the same decontX and Scrublet quality control pipeline before merging with the patch-seq object. The final integrated object included SCβ-cells, primary endocrine cells, and S6D1 cells across all studies.

Three normalized data representations were derived from this object for different analytical purposes. Log-normalized counts (RNA data slot) were used for transcript–electrophysiology correlation analysis, as Spearman correlations are robust to the distributional assumptions of this normalization. For within-population correlation analyses shown in **Fig. 4A,B**, SCTransform Pearson residuals (SCT scale.data slot) were used instead, which are variance-stabilized across cells and appropriate for visualizing relative expression differences within the SCβ-cell population. SCTransform Pearson residuals (SCT scale.data slot) were also used for MOFA2 unsupervised integration to satisfy the Gaussian likelihood assumption of that framework. SCTransform-normalized expression aggregated to pseudobulk level and normalized by group size (SCT data slot, summed then divided by cell count per Differentiation × cell type group) was used for DIABLO supervised multi-omics integration. SCENIC regulon AUC values were stored in a separate AUC assay and used directly for co-activity network analysis, MOFA2, and DIABLO.

#### Differential Expression

Differential expression between D30 and D50 SCβ-cells was assessed using a bootstrapped pseudobulk approach. SCβ-cells were subsetted into D30 and D50 groups (above), and raw RNA counts were extracted. Genes detected in at least 10% of SCβ-cells were retained for analysis. Cells were then aggregated into pseudobulk samples by differentiation and cell stage. To estimate the stability of differential expression, pseudobulk samples from each group were resampled with replacement for 100 bootstrap iterations. At each iteration, differential expression was tested using a generalized Poisson model with cell stage as the design variable and D30 SCβ-cells as the reference group. The contrast tested genes enriched or depleted in D50 SCβ-cells relative to D30 SCβ-cells. Bootstrap results were summarized for each gene by calculating the mean and standard deviation of log2 fold change, adjusted p value, signed test statistic, and gene deviance across iterations. Genes were considered stable if they were significant in at least 90% of bootstrap iterations and had a mean adjusted p value ≤ 0.05. Stable genes were used for downstream visualization, including volcano plots highlighting the strongest upregulated and downregulated genes.

Differential expression between SCβ- and primary β-cells was also performed on pseudobulked counts, aggregated by Differentiation × Study × cell type, using a negative binomial generalized linear model implemented in glmGamPoi. To stabilize estimates across the heterogeneous multi-study dataset, 100 bootstrap resamples were generated by sampling pseudobulk samples with replacement within each cell type, and a model was fit to each resample. Genes were summarized by mean log fold change, directional consistency (fraction of bootstraps with positive log fold change), and a signed effect size statistic. Leave-one-study-out (LOSO) sensitivity analysis was performed by iteratively excluding each primary β-cell study, repeating the bootstrap procedure, and computing the relative stability of effect size estimates across exclusions. A within-study sensitivity analysis using only the smaller primary β-cell and SCβ study was also performed. Genes were triaged into four tiers based on effect size magnitude and LOSO stability: Tier1A (strong effect, stable across studies, supported by SCβ-cell sensitivity analysis), Tier1B (strong, stable, SCβ data unavailable), Tier2 (strong but sensitive to study exclusion), Tier3 (weak but stable), and Tier4 (weak and unstable). Tier1A genes were used for labeling, interpretation, and downstream feature selection (**Supplementary Table 1**).

#### Transcript-Electrophysiology Correlations

For within-population correlation analyses (**Fig. 4A,B**), bootstrapped Spearman correlations were computed within the SCβ population only, using SCTransform Pearson residuals (SCT scale.data). Genes detected in ≥5% of cells were retained. Correlations were computed across 100 bootstrap resamples (cells sampled with replacement), and summarized as mean correlation, 95% bootstrap confidence intervals, and empirical pseudo-p values based on zero-crossing frequency, FDR-adjusted within each electrophysiology parameter. Current amplitudes were sign-flipped so that more positive values indicate greater current magnitude. This analysis characterizes transcript–electrophysiology relationships within SCβ-cells.

To identify divergent transcript–electrophysiology relationships between SCβs and primary β-cells (**Fig. 4C-G**), correlations were recomputed using log-normalized RNA expression values. Genes were retained if detected in ≥10% of cells in both populations and if their standard deviation exceeded the 10th percentile in at least one population, yielding approximately 10,000 genes. For primary β-cells, Spearman correlations were computed within each of five independent studies across 200 bootstrap resamples, each randomly drawing one of 50 imputed electrophysiology datasets to propagate imputation uncertainty. Study-specific Fisher z-transformed estimates were combined using DerSimonian-Laird random-effects meta-analysis, yielding a pooled z-estimate, standard error, and I² per gene-parameter pair. For SCβ-cells, bootstrapped Spearman correlations were computed on resampled cells across the same 200 iterations. The difference in Fisher z-transformed correlations (Δz = z_SCβ − z_β) was used as the primary divergence metric. High-confidence associations were defined by sign stability ≥0.9 and 95% CI width ≤0.3. Gene set enrichment analysis was performed using fgsea on Δz-ranked gene lists against Hallmark gene sets.

#### Multiple Imputations for Missing Electrophysiology Data

Missing electrophysiology values were handled using multiple imputation by chained equations (MICE) implemented in the mice R package, with m = 50 imputed datasets. Imputation models were tailored to the measurement characteristics of each parameter type. Exocytosis parameters were imputed using a two-part model: logistic regression for presence or absence of an exocytotic response, followed by log-transformed predictive mean matching for response magnitude, conditional on presence. Negative exocytosis values, reflecting cells with no measurable exocytotic response, were retained as real observations rather than treated as missing, and were reconstructed as small negative values after imputation. Inward current parameters (Ca^2+^ and Na^+^ currents) were imputed as magnitudes using a censored Gaussian model via Tobit regression, then restored to signed negative values. Voltage parameters were imputed using midastouch, except for voltage of peak Na^+^ current which was treated as an ordered factor and imputed with ordered logistic regression. Predictors for all imputation models included cell type, study, donor, cell size, glucose concentration, diabetes status, and patcher identity. Biological sign constraints were enforced post-imputation, and cell-size-normalized parameters were recomputed from reconstructed signed values for each completed dataset. Imputation quality was assessed by comparing observed and imputed distributions using density plots and Kolmogorov-Smirnov tests. Downstream analyses randomly sampled one imputed dataset per bootstrap iteration to propagate imputation uncertainty through all statistical estimates.

#### SCENIC regulon inference and co-activity networks

Gene regulatory network (GRN) inference was performed using pySCENIC on the quality-controlled scRNA-seq dataset. To obtain a robust consensus GRN, the inference step (pyscenic grn) was run 100 times with different random seeds using SLURM job arrays, and transcription factor-target edges were retained only if present in 90 or more of 100 runs. Edge importance scores were averaged across runs to yield a consensus GRN, which was used as input for the cisTarget (ctx) and AUCell steps to generate regulon definitions and per-cell regulon activity scores (AUC values).

Inferred undirected co-activity network plots across primary β-cell, SCβ-cells, and S6D1 cells were generated by computing pairwise Spearman correlations on regulon AUC values. Edges were retained at ρ greater or equal to 0.55 and only if present in greater or equal to 50/100 bootstraps (cells resampled with replacement) to limit sampling noise/sparsity. Graphs were constructed from symmetric pairwise Spearman correlations; edge directionality was not biologically informative in this context. Regulons were colored according to differential network influence between SCβ-cells and primary β-cells. We further organized these into biologically relevant programs.

We then computed PageRank centrality (damping factor = 0.85) per transcription factor on correlation-derived graphs (ρ>0.30) across 100 bootstraps (of 80% cell resampling). Δ Influence = mean(PageRank_β_) - mean(PageRank_SCβ_), and we kept the top 10 by absolute Δ Influence for downstream analysis. For directed GRN subnetwork visualization, influential transcription factor-target edges from the consensus GRN (importance scores above the 95th percentile of the consensus GRN importance distribution) formed a directed subnetwork for the top 13 transcription factors by absolute Δ Influence. Networks were visualized using a stress-minimization layout. Differential activity/expression was detected by BH-FDR < 0.05 (node colours reflect differential expression or influence direction). These analyses were used to identify highly influential transcription factors. For supplementary themed subnetwork figures, target genes were functionally annotated using Gene Ontology Biological Process enrichment (clusterProfiler) with semantic similarity reduction via rrvgo (threshold = 0.7).

#### DIABLO analysis

Supervised multi-omics integration was performed using DIABLO (block sPLS-DA; mixOmics)^22^ on pseudobulked data aggregated by Differentiation/Donor × cell type. Three data blocks were used: (i) SCT-normalized mRNA expression, normalized by group size, scaled, and filtered to remove near-zero variance features and LOSO-unstable genes; (ii) regulon AUC values from SCENIC, aggregated and normalized by group size; and (iii) cell-size-normalized electrophysiology parameters, with one imputed dataset used per model run. The full design matrix was used (all pairwise block correlations weighted equally). The optimal number of components was determined by 10-fold cross-validation repeated 100 times, selecting by balanced error rate (BER) using centroids distance. The number of features per block was tuned using the same cross-validation framework across a grid of candidate values.

To obtain robust consensus features, DIABLO was run independently on each of the 50 imputed electrophysiology datasets, yielding 50 models. Loadings were sign-aligned across models using Procrustes-style correlation with a reference model. Features were considered consensus if selected in ≥80% of models with ≥90% sign consistency across models. Cross-block Pearson correlations between selected features were averaged across models to yield a consensus correlation matrix. Features present in fewer than 20% of models or matching unannotated transcript patterns were excluded. The consensus cross-block correlation matrix was visualized as a heatmap of ephys × transcriptomic feature correlations, with row and column annotations showing standardized effect sizes from linear mixed models (SCβ vs. primary β-cells, Differentiation as random effect; see SCENIC regulon analysis below) using a shared colour scale across both annotation tracks.

Differential activity of SCENIC regulons, mRNA features, and electrophysiology parameters selected by DIABLO was assessed using linear mixed models (lme4),^45^ with cell type (SCβ-cells vs. primary β-cells) as a fixed effect and differentiation batch or donor as a random effect, restricted to the current study to avoid cross-study confounding. Effect sizes were standardised by dividing the fixed effect estimate by the standard deviation of the outcome variable, yielding a metric comparable to Cohen’s d. Models showing singular fits were refitted as fixed-effects linear models. P-values were adjusted using the Benjamini-Hochberg procedure. Standardised effect sizes from these models were used to annotate the DIABLO cross-block correlation heatmap, with a shared color scale across row (electrophysiology) and column (regulon and mRNA) annotation tracks to enable direct comparison of effect magnitudes across data modalities.)

#### MOFA2 Analysis

Unsupervised multi-omics factor analysis was performed using MOFA2^23^ independently on each of 50 imputed electrophysiology datasets. Three views were provided: (i) SCT-normalized mRNA expression represented as Pearson residuals from the SCTransform regularized negative binomial regression (SCT scale.data slot), which are approximately normally distributed and variance-stabilized, consistent with the Gaussian likelihood assumption of MOFA2; features were filtered to the top 4,000 genes by variance after removing LOSO-unstable genes and ribosomal protein genes; (ii) scaled regulon AUC values from SCENIC, filtered to remove near-zero variance features; and (iii) sign-corrected scaled electrophysiology parameters, with S6D1 cells assigned NA values (handled natively by MOFA2 as missing views). Each model was trained with 10 factors using slow convergence mode, with seeds varied per imputation run.

To obtain a consensus separation factor across imputation runs, each model’s factors were evaluated for their ability to separate SCβ-cells and S6D1 cells from primary endocrine cells, quantified as the standardized mean difference in factor scores between groups. The factor with maximum absolute separation was selected per model, and sign-aligned across models by correlation with a reference model. Consensus factor scores were computed as the mean across 50 sign-aligned models and used for downstream visualization and analysis. Feature weights were summarised across models, retaining features with ≥90% sign consistency; the top 25 features per view by mean absolute weight are reported.

For visualization, consensus factor scores were embedded in two dimensions using UMAP (n_neighbors = 30, min_dist = 0.3, Euclidean metric). Consensus factor scores per cell type were visualized as violin plots. The diverging lollipop plot shows mean feature weights ± SD across 50 models for the top features from the electrophysiology and SCENIC views, with positive loadings indicating association with the SCβ/S6D1 state and negative loadings indicating association with primary endocrine cell identity.

#### Network control energy and transcription factor prioritization

Transcription factor-target regulatory dynamics were modeled as a linear time-invariant system ẋ = Ax + Bu, where A is the row-normalized adjacency matrix derived from the consensus GRN, B is an input matrix selecting TF nodes, and x represents MOFA2 consensus factor projections. Rather than computing true minimum control energy via the controllability Gramian, input energy was estimated as the squared L2 norm of each transcription factors’s predicted input when the system is driven along the trajectory from the mean SCβ-cell MOFA2 factor state toward the mean primary β-cell MOFA2 factor state via forced response simulation over T = 10 time steps. This approach identifies relative rankings of candidate regulators but does not provide theoretically optimal energy estimates, and assumes a linear approximation of gene regulatory network dynamics. Transcription factor rankings were stabilized across 100 bootstrap resamplings of cells (80% resampling fraction). Each transcription factor was characterized by (i) Δ influence from co-activity network analysis, and (ii) predicted input energy from the forced response simulation.

## Data and Code Availability

Data and code will be available at the time of publication.

## Declaration of generative AI and AI-assisted technologies in the writing process

During the preparation of this work, the author(s) used OpenAI Codex and ChatGPT to assist with grammar and wording, code planning/review, and script cleanup. Claude (Anthropic, claude.ai) was used to refactor analytical code into reusable functions and loops, to annotate and document R and Python scripts for reproducibility, and to assist in drafting the methods section based on author-provided code and analytical decisions. After using this tool or service, the author(s) reviewed and edited the content as needed and take(s) full responsibility for the content of the publication.

## Acknowledgements

The University of Alberta acknowledges it is located on Treaty 6 territory and honours the histories, languages, and cultures of First Nations, Métis, Inuit, and all First Peoples of Canada, whose presence continues to enrich our community. We gratefully acknowledge all donor families for their generous contributions to diabetes and transplantation research, and thank Give Life Alberta (AHS), Trillium Gift of Life Network (TGLN), BC Transplant, Quebec Transplant, and other Canadian organ procurement organizations for pancreas procurement.

We thank James Johnson (University of British Columbia) for helpful comments. We acknowledge the Faculty of Medicine and Dentistry Cell Imaging Core and Facility for TEM. This research was enabled in part by support provided by Prairies DRI and the Digital Research Alliance of Canada (alliancecan.ca).

## Author Contributions

J.M. performed experiments, analysis, and wrote the manuscript. C.E. performed analysis and wrote the manuscript. A.F.S. performed experiments. N.S. performed experiments. S.S performed experiments. F.C.L provided cells and edited the manuscript. P.E.M supervised the work and edited the manuscript. All authors have reviewed and approved the manuscript.

---

**Supplementary Figure S1:**
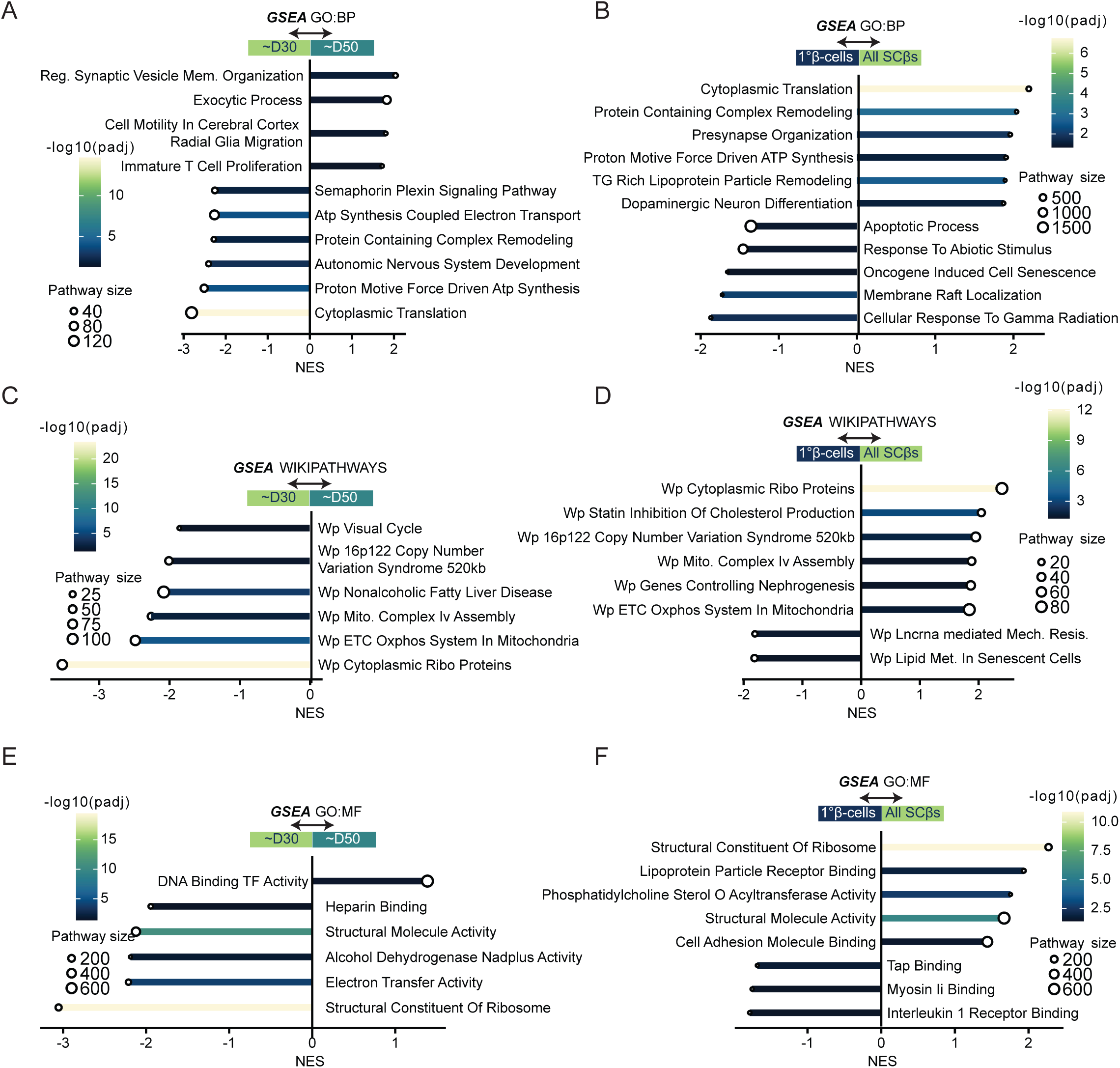
Supplementary Gene Set Enrichments. Gene set enrichment analysis (GSEA) was performed using GO Biological Process (GO:BP, Panels A-B), GO Molecular Function (GO:MF, Panels C-D), and WikiPathways (Panels E-F) gene sets on ranked differential expression results for D30 versus D50 SCβ cells (*left side of page*) and primary β-cells versus SCβ cells (*right side of page*). Enriched pathways are shown as normalized enrichment scores (NES), with significance determined by adjusted P values/FDR as indicated.

**Supplementary Figure S2:**
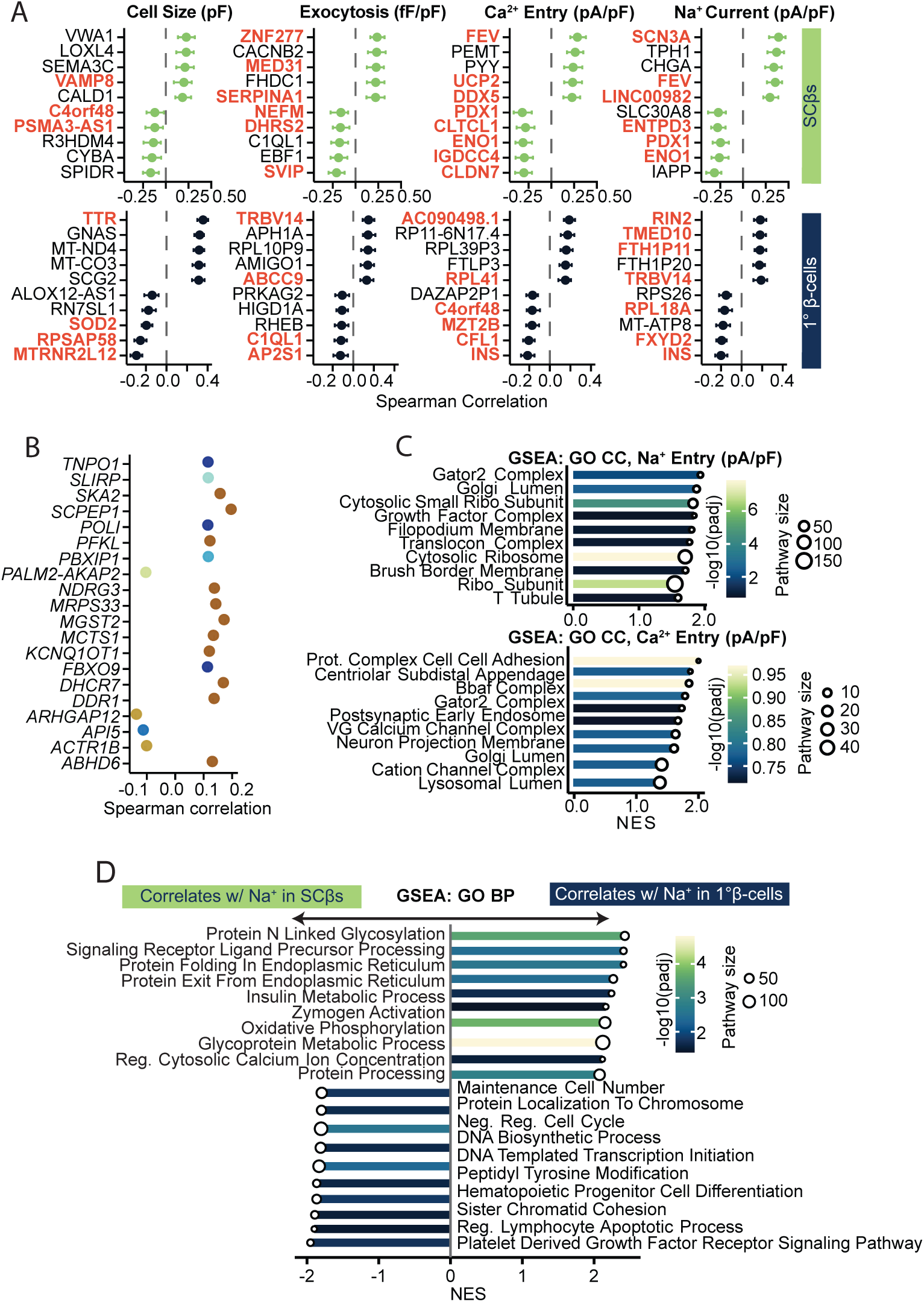
Supplementary Transcript - Electrophysiology Relationships. (A) Divergent transcript-electrophysiology relationships between primary and SCβ-cells. Error bars represent the confidence interval of bootstrapped values. (B) Convergent transcript-electrophysiology relationships identified by bootstrapped correlation analysis and random-effects meta-analysis, highlighting genes with consistent direction and magnitude of association across primary and SCβ-cells. (C) Functional enrichment of convergent genes, showing pathways related to cell-cell adhesion, centriolar subdistal appendages, BAF complex, postsynaptic early endosomes, voltage-gated calcium channel complexes and Golgi lumen. GSEA of genes ranked by divergence effect size for sodium current. Bars show normalized enrichment score (NES), color encodes -log10(adjusted p) and point size indicates pathway size.

**Supplementary Figure S3:**
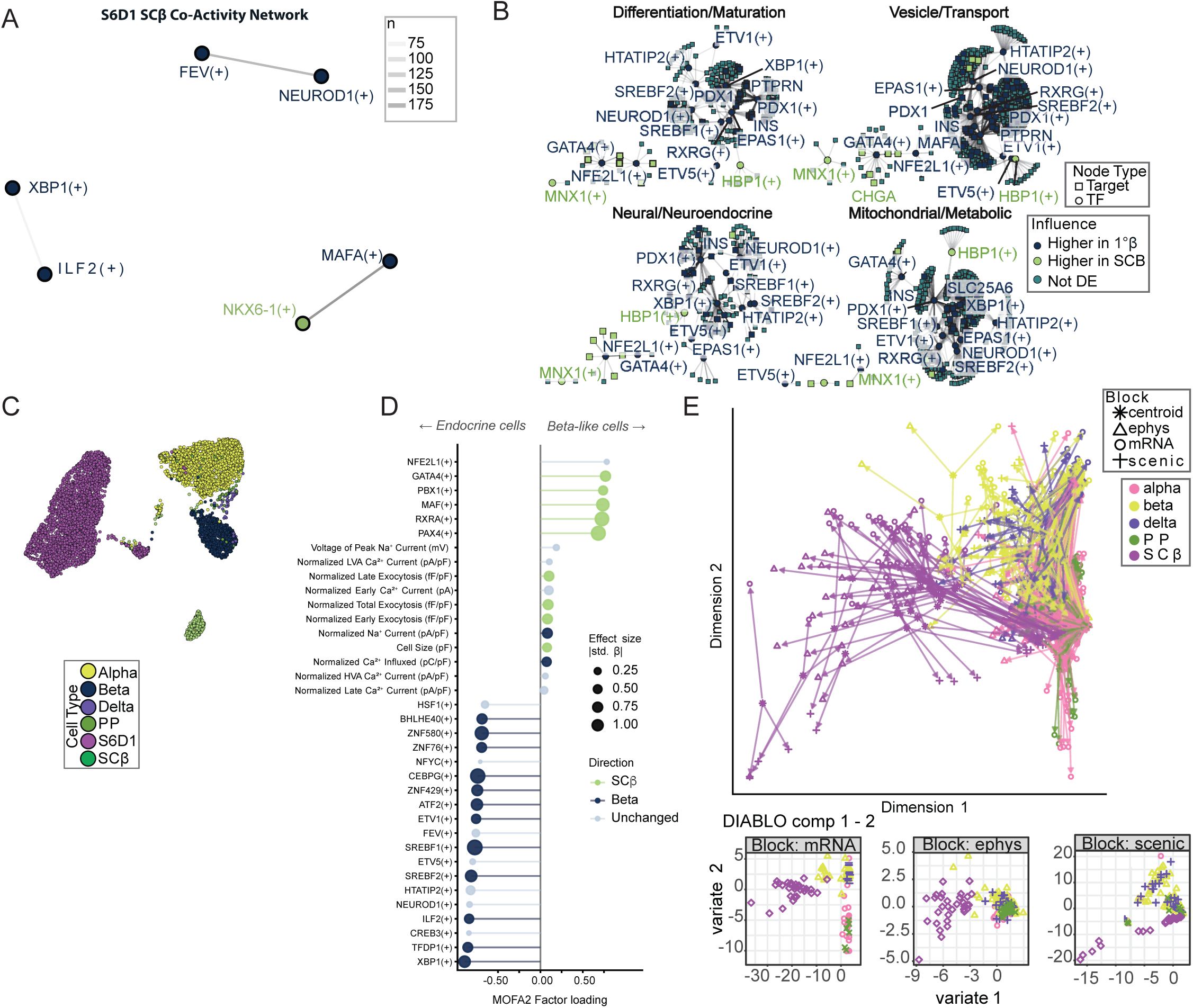
Supplementary Regulatory Network Analyses. (A) Bootstrapped co-activity networks derived from SCENIC AUC values for S6D1 stem cells.. Edges represent reproducible regulon co-activity across bootstrap iterations (present in ≥50% of iterations). Nodes are colored according to condition-specific upregulation. (B) Top 10 transcription factors (TFs) ranked by bootstrapped delta influence between SCβ-cells and primary β-cells, together with their top target genes. Nodes are shaped by type (TF or target) and colored by direction of differential expression. TFs with greater influence in SCβ-cells are broadly rewired and often regulate targets that are also upregulated in SCβ-cells, suggesting active roles in altered gene regulatory programs. (C) UMAP embedding of consensus MOFA2 factor scores (mean across 50 imputation runs), computed using all 10 factors (n_neighbors = 30, min_dist = 0.3, Euclidean metric), colored by cell type. Primary endocrine cell types form distinct clusters while SCβ-cells occupy a separate region consistent with a transcriptionally coherent but divergent identity. (D) Consensus MOFA2 primary separation factor loadings for electrophysiology parameters (circles) and SCENIC regulon AUC values (triangles). Points show mean loading across 50 imputation runs; only features with ≥90% sign consistency across runs are shown, representing the top 25 features per view by mean absolute loading. Positive loadings indicate association with the SCβ/S6D1 state; negative loadings indicate association with primary endocrine cell identity. Current amplitude parameters were sign-flipped prior to analysis so that more positive values indicate greater electrophysiological activity. (E) Representative DIABLO model diagnostics from a single imputation run. Top: arrow plot (plotArrow) showing the displacement of each sample between its block-specific projections on components 1 and 2; shorter arrows indicate greater cross-block agreement for that sample. Bottom: block-specific sample scores on components 1 and 2 (plotIndiv) for mRNA, electrophysiology, and SCENIC views separately, colored by cell type. SCβ-cells (pink) are clearly separated from primary endocrine cell types across all three data blocks.

## SUPPLEMENTARY TABLES

**Suppl Table 1:** Differential gene expression in human primary β-cells versus SCβ-cells.

**Suppl Table 2:** Correlation analysis between gene transcripts and electrophysiological features.

**Suppl Table 3:** Divergent gene regulatory networks (regulons) and their downstream targets.

**Suppl Table 4:** Latent factor loadings from multi-omics analysis (MOFA2).

**Suppl Table 5:** Control theory ranking and prioritization of transcription factors.

**Suppl Table 6:** Gene expression changes in SCβ-cells following U18666A treatment.

